# Evolutionary patterns and heterogeneity of Dengue Virus serotypes in Pakistan

**DOI:** 10.1101/2023.09.28.559885

**Authors:** Zilwa Mumtaz, Rashid Saif, Muhammad Zubair Yousaf

## Abstract

Comprehensive and systematic examination of Dengue virus (DENV) evolution is essential in the context of Pakistan as the virus presents a significant public health challenge with the ability to adapt and evolve. To shed light on intricate evolutionary patterns of all four DENV serotypes, we analyzed complete genome sequences (n=43) and envelope (E) gene sequences (n=44) of all four DENV serotypes collected in Pakistan from 1994 to 2023 providing a holistic view of their genetic evolution. Our findings revealed that all four serotypes of DENV co-circulate in Pakistan with a close evolutionary relationship between DENV-1 and DENV-3. Genetically distinct serotypes DENV-2 and DENV-4 indicate that DENV–4 stands out as the most genetically different, while DENV-2 exhibits greater complexity due to the presence of multiple genotypes and the possibility of temporal fluctuations in genotype prevalence. Selective pressure analysis in Envelope (E) gene revealed heterogeneity among sequences (n=44) highlighting 46 codons in the genome experiencing selective pressure, characterized by a bias towards balancing selection indicating genetic stability of the virus. Furthermore, our study suggested an intriguing evolutionary shift of DENV-4 towards the DENV-2 clade, potentially influenced by antibodies with cross-reactivity to multiple serotypes providing a critical insight into the complex factors shaping DENV evolution and contributing to the emergence of new serotype.

**Author Summary:** The emergence of the fifth serotype of dengue virus heightened our interest in investigating its presence in Pakistan. In our quest to understand the evolving landscape of dengue in Pakistan, we conducted a comprehensive analysis, comparing whole genome sequences and E gene sequences. Notably, we focused on the E gene recognized as the most mutable component and a key determinant of dengue’s virulence. The phylogenetic analysis unveiled fascinating findings, demonstrating a strong genetic affinity between serotypes 1 and 3. Substantially signifying its implications for vaccine development and understanding of cross-immunity dynamics within serotypes. We delved into the genetic dynamics of dengue by subjecting the whole genome of DENV and E gene to neutrality tests. The outcomes of these tests unveiled a critical aspect of dengue virus evolution: the genome is not evolving neutrally. Instead, the E gene experiences selective pressure, indicating a bias towards balancing selection. The finding underscores the complex interplay of factors shaping the genetic diversity of dengue in Pakistan and provides valuable insights into the virus’s adaptive strategies.

## Introduction

Dengue infection is prevalent among humans and is primarily found in regions characterized by the tropical and sub-tropical climates. It is estimated that there are approximately 400 million new infections each year on a global scale. The illness caused by dengue fever encompasses a spectrum of manifestations ranging from an acute febrile condition known as dengue fever to dengue hemorrhagic fever or dengue shock syndrome which is a more severe and potentially life-threatening form. The single stranded positive RNA virus behind dengue infection belongs to virus Flaviviridae having a genome size of 11 kb and encodes a polyprotein with a mutation rate of more than 100 times greater than mutation rates of DNA genomes [1].

Neutral theory of molecular evolution states the genetic differences between species and within a population are two sides of the same coin. Because, a large number of evolutionary substitutions that manifest as species differences are mostly neutral, meaning that a specie could accept these without having a substantial adverse impact on its capacity to survive. These variations are termed as evolutionary permissible variations. It is possible to deduce evolutionary permissibility of every potential variant by using multispecies sequence alignment and sophisticated statistical techniques on the evolutionary tree of species. However, evolutionary forbidden variants in the genome may impart an unappreciated role in protection against disease and evolution of new genomes from previous ones. The infection of DENV begins with host cell adhesion, entrance and virus envelope fusion with cellular endosomal membranes carried out by the Class II fusion protein known as the flavivirus E protein. E gene is regarded as a molecular marker of DENV pathogenicity because E protein’s adhesion to the target cell containing heparin sulfate is necessary to initiate DENV infection. The genomic sequence of E gene also differs in pathogenic and non-pathogenic strains. Therefore, the antibodies that successfully inactivate one serotype may not work against another due to the structural differences in the E gene [2].

Our study represents a pioneering effort in Pakistan, as it provides the first comprehensive examination of the dengue virus in the region, focusing on its evolutionary landscape. By assembling a dataset comprising a total of 43 complete DENV genomes from serotype 1 to 4 and 46 E gene sequences, we have created the extensive collection of DENV data from Pakistan to date. The phylogenetic analysis unveiled a compelling relationship between DENV-2 and DENV-4. However, it is crucial to recognize the co-circulation of multiple DENV serotypes coupled with heightened human activity, elevated the risk of genetic changes that contribute to the diversity within virus populations. The study also highlights the role of natural selection and genetic bottlenecks in the emergence of new dengue virus serotypes, further emphasizing the significance of ongoing monitoring and research in combating this significant public health threat. DENV Envelope (E) gene selection analysis indicated the existence of diversifying and purifying selection along with two recombination breakpoints that indicates a departure from strict neutrality and the presence of selective forces acting upon viral sequences.

## Results

### DENV Evolutionary Analysis

Multiple sequence alignment of whole genome sequences testified the 100% conservation of 5ꞌ and 3ꞌ cyclization sequences in all 43 sequences.Fig 1 represents the GC content of DENV-2 reference genome and its complete genomic representation.

**Fig 1.**
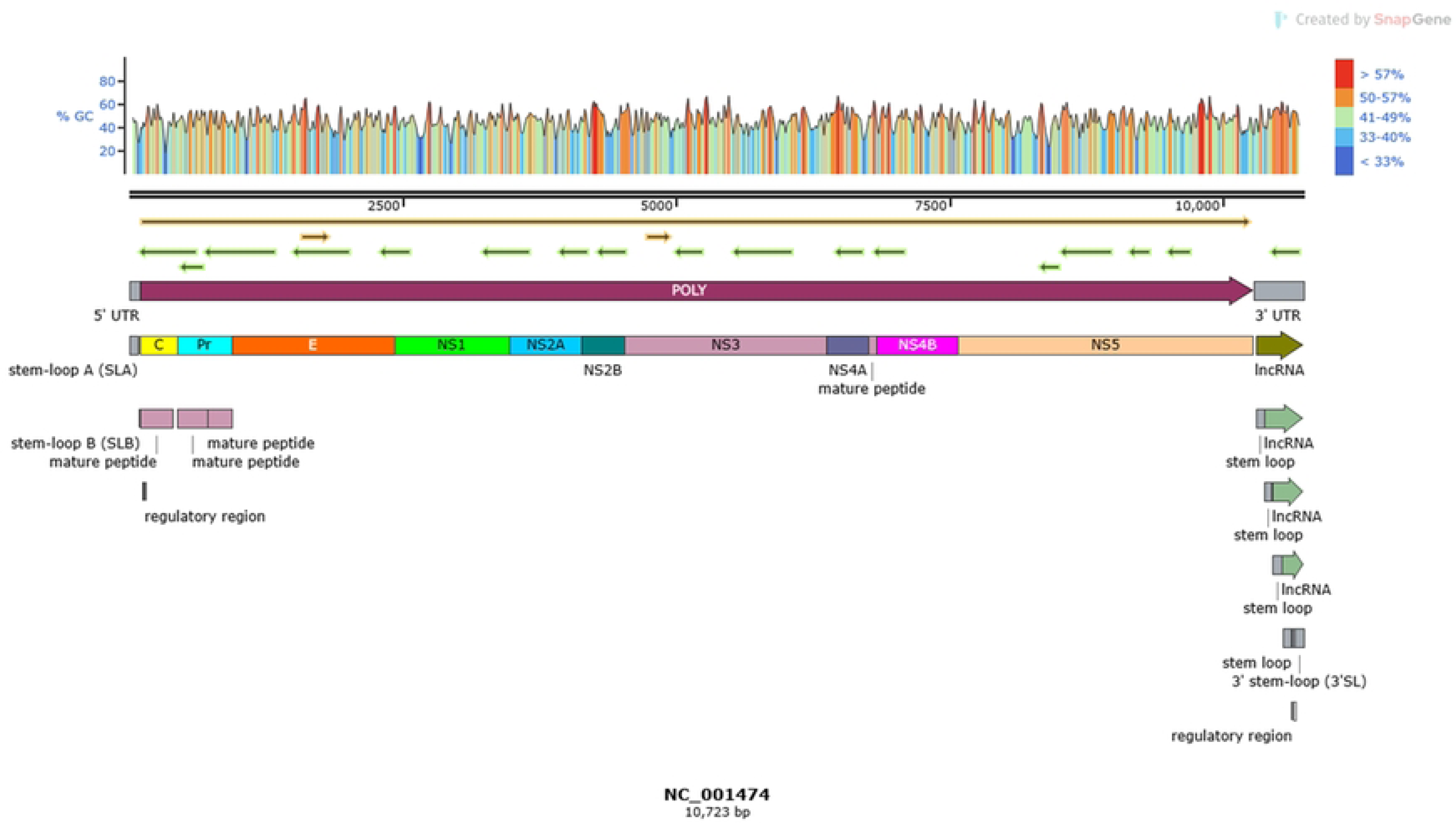
Complete Genomic representation of DENV-2 reference genome with accession no. (NC_001474.2). Percentage of GC content is shown in the entire genome where red represents nucleotides with more than 57 GC content. Orange color shows 50-57%, green color corresponds to 41-49%, light blue color shows 33-40% and dark blue color represents less than 33% GC content in nucleotides. (created in SnapGene).

Whole genome of DENV translates into a single Polyprotein under one ORF. The polyprotein further divides into 3 structural and 7 non-structural proteins. The region from 97-438 bp encodes anchored capsid protein ancC, 439-936 encodes membrane glycoprotein precursor prM, 712-936 encodes membrane glycoprotein M, 937-2421 encodes envelope protein E, 2422-3477 encode nonstructural protein NS1, 3478-4131 encodes nonstructural protein NS2A, 4132-4521 encodes nonstructural protein NS2B, 4522-6375 encodes nonstructural protein NS3, 6376-6756 encode nonstructural protein NS4A, 6826-7569 encode nonstructural protein NS4B and 7570-10269 encodes nonstructural protein NS5.

Phylogenetic analysis of 43 complete DENV genome sequences showed a compelling pattern among serotypes. The bootstrap values are calculated from 1000 replicates and the percentage supporting value is represented on nodes. The scale bar represents 0.050 substitutions per site.

DENV-1 and DENV-3 shared one main clade and subdivided into two sub-clades, while DENV-2 and DENV-4 shared another main clade and divided into two sub-clades as shown in Fig 2. Evolutionary relatedness of dengue virus serotypes and genotype prevalence is shown in S1 Fig. DENV-3 sequences clustered into Genotype-III, while DENV-1 along with the sole DENV-4 sequence, exhibited Genotype-I. DENV-2 showed intriguing diversity, the 10 sequences of DENV-2 collected in 2022 shared Genotype-I suggesting its prevalence. whereas, two older DENV-2 sequences shared Genotype I and IV. (Sylvatic). Whereas, 12 sequences from DENV-2 clustered into an unknown genotype.

**Fig 2.**
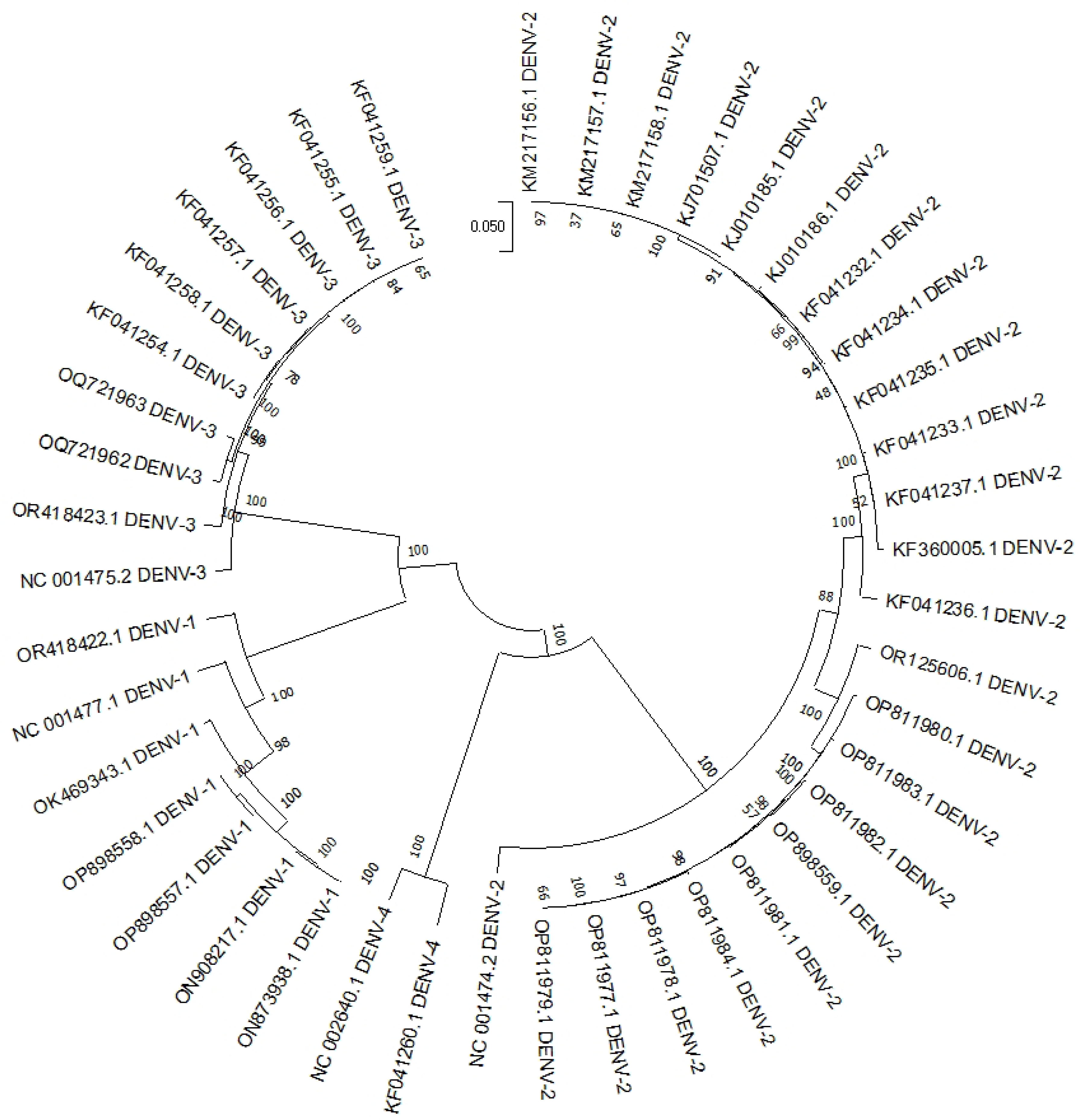
Phylogenetic analysis of 43 complete genome sequences of DENV-1, DENV-2, DENV-3 and DENV-4 serotypes. Maximum likelihood (ML) tree was constructed with bootstrap value (1000) using the GTR+G+ I model.

### Departure from neutrality

The Fisher’s Exact test of neutrality revealed a significant departure from strict neutrality, with likelihood values supporting the alternative hypothesis of positive selection for sequence pairs. Similarly, the Codon based Z test indicated significant departures from neutrality with P values less than 0.05, underscoring selective pressure acting on these sequences. Furthermore, the examination of non-synonymous and synonymous substitutions per site unveiled insights into specific selective pressure at play. The employment of the Discrete Gamma Distribution, with a shape parameter of 0.3926, effectively categorized evolutionary rates into five distinct groups. These rates ranged from 0.01 to 3.50 substitutions per site, providing a detailed view of the varying evolutionary dynamics across different sequence positions. Regarding nucleotide composition, the frequencies exhibited by sequences were A (32.48%), T/U (21.19%), C (20.61%) and G (25.71%). These percentages delineate the compositional makeup of the genetic sequences. R value of 1.94 obtained in case of Transition/Transversion bias. Tajima’s D test, conducted on 43 complete genome sequences with a total of 5593 sites, further illuminates the dataset. A noteworthy 51.7% of the sites showed genetic variations (S/n ≈ 0.517) indicating a substantial genetic diversity. Nucleotide diversity (π ≈0.20) reflected a moderate level of genetic variation, and Tajima’s D test statistic was 2.545 as depicted in Table 1, further underscoring the deviation from neutrality in the dataset.

**Table 1.** Regions under genomic selection pressure measured by Tajima’s D test.

| *m* | *S* | *p<sub>s</sub>* | $\Theta$ | $\pi$ | *D* |
| --- | --- | --- | --- | --- | --- |
| 43 | 5593 | 0.517056 | 0.119502 | 0.200995 | 2.545272 |

**Table 2.**
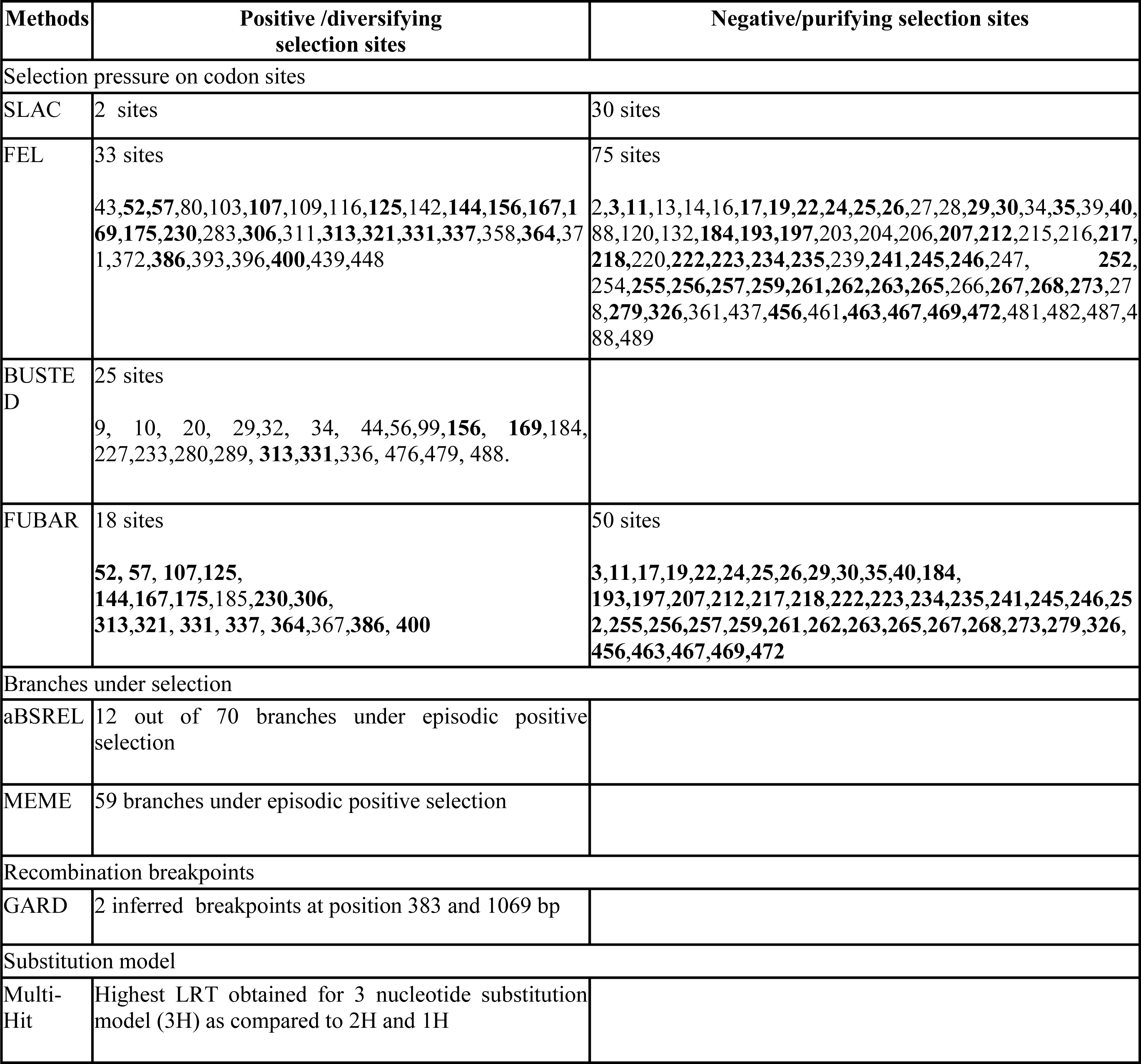
Selection Pressure analysis tests results and their comparison. The codons confirmed by at least two Tests are shown as Bold numbers.

Here, *m* represents the number of sequences, *S* refers to segregating sites, *π* represents nucleotide diversity and *D* is the Tajima test statistic.

### Envelope gene evolutionary analysis

E gene phylogenetic analysis suggested a common ancestor and close relationship between DENV-1 and DENV-3 forming a distinct clade, while DENV-2 and DENV-4 are more genetically divergent from other two serotypes indicated by their higher genetic distance. Moreover, DENV-4 with the highest genetic distance appeared to be the most genetically distinct serotype among others as shown in Fig 3.

**Figure 3.**
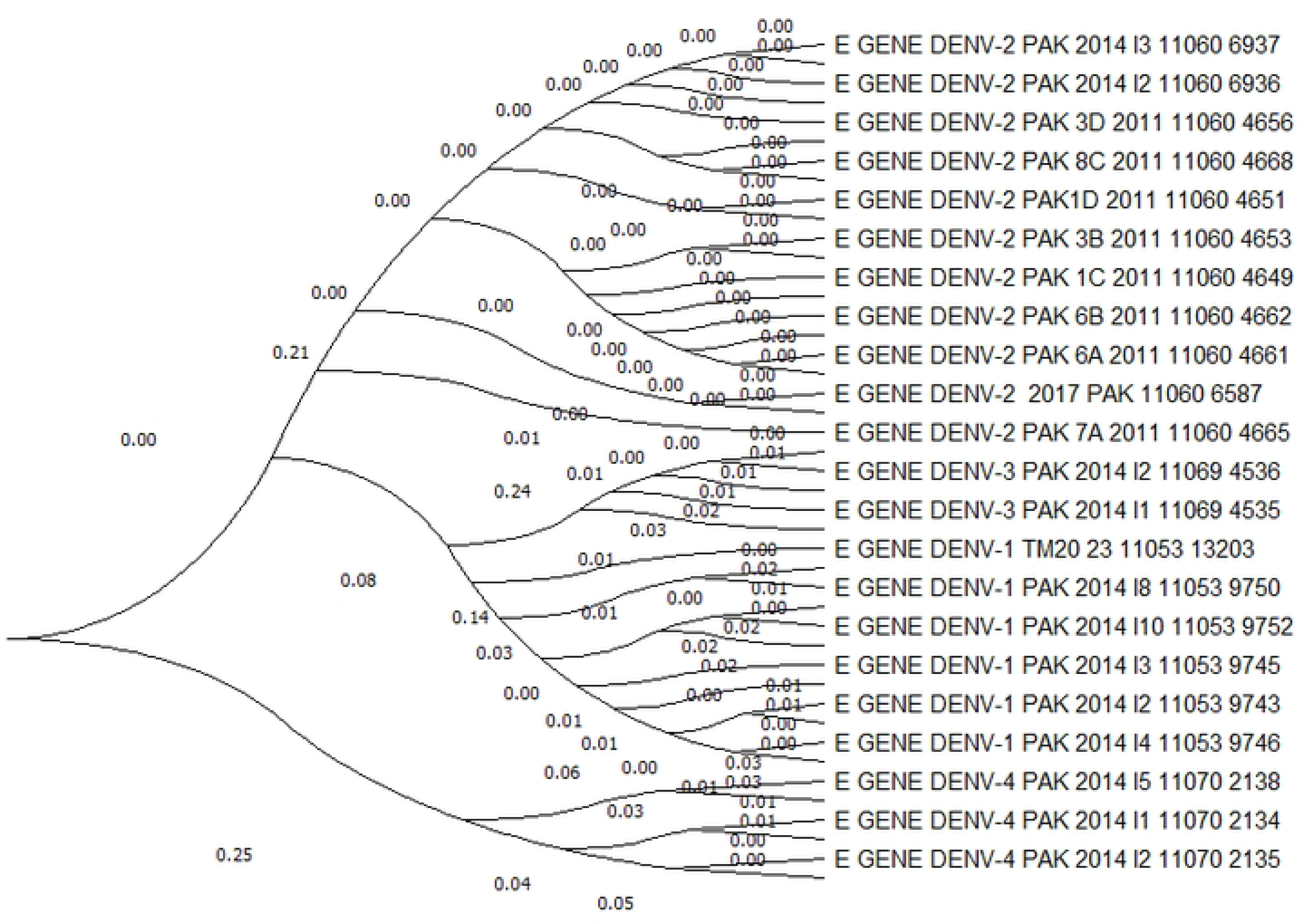
Evolutionary analysis of Dengue virus serotypes (DENV-1, DENV-2, DENV-3 and DENV-4) based on envelope gene sequences. DENV-1 and DENV-3 share a common clade, indicating a closer evolutionary connection. The tree was constructed using Maximum Likelihood with Tamura-Nei model, and the displayed tree represents the highest log likelihood (−9153.31). Branch lengths indicate substitutions per site.

### Inferred selection pressure

2 sites were selected in SLAC that were under diversifying selection and 30 under purifying selection with p-value ≤ 0.1. The nucleotide GTR model was selected with the highest log L value to determine rates of synonymous and nonsynonymous mutations and their effect on amino acids as shown in Fig 4.

**Fig 4.**
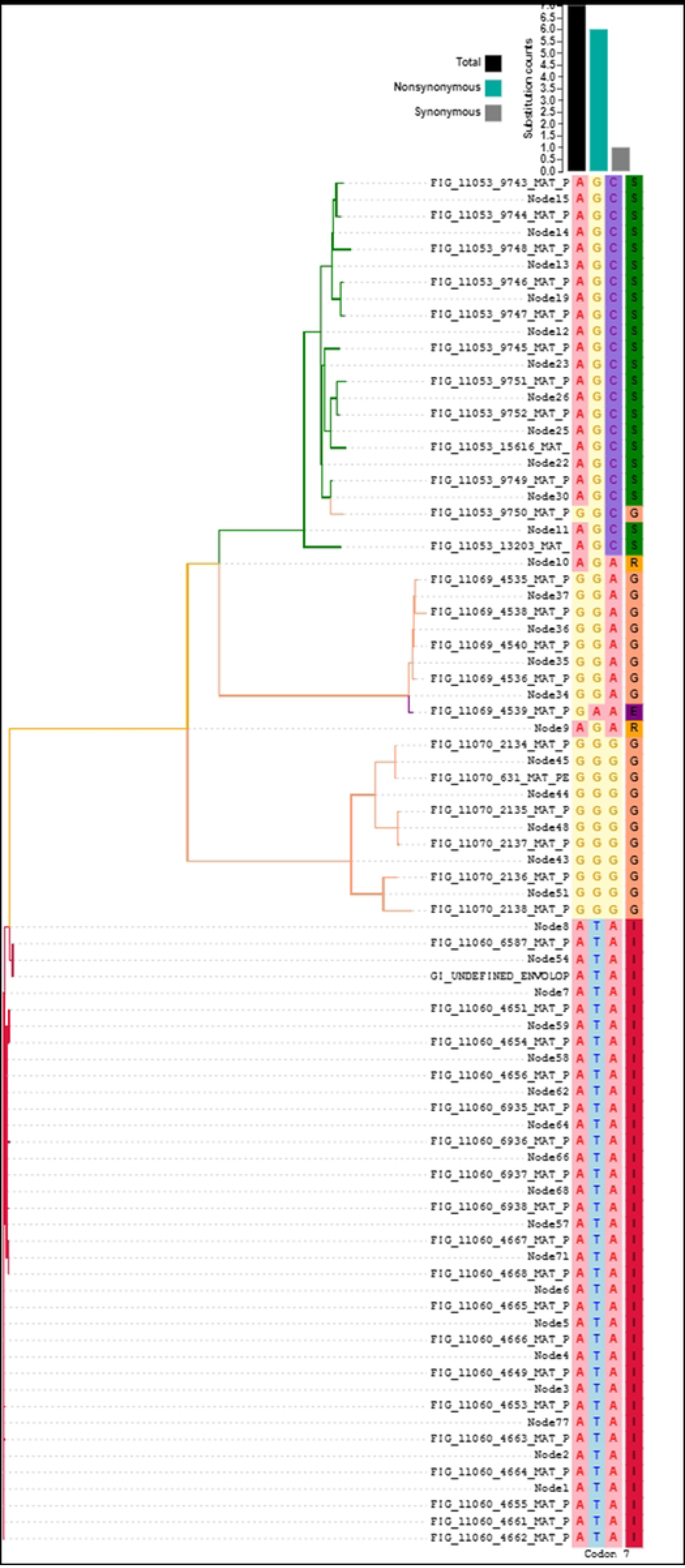
Nonsynonymous and synonymous mutations in codons no. 7 affecting amino acid composition. The number of non-synonymous mutations are more as compared to synonymous mutations.

The FEL analysis identified 75 sites that were under purifying selection and 33 under diversifying selection at p≤0.1. Fig 5 shows a dense rate plot in which alpha and beta substitutions are shown in codons. The peaks in the plot correspond to sites within the E gene indicating synonymous substitutions (α) and non-synonymous substitutions.

**Fig 5.**
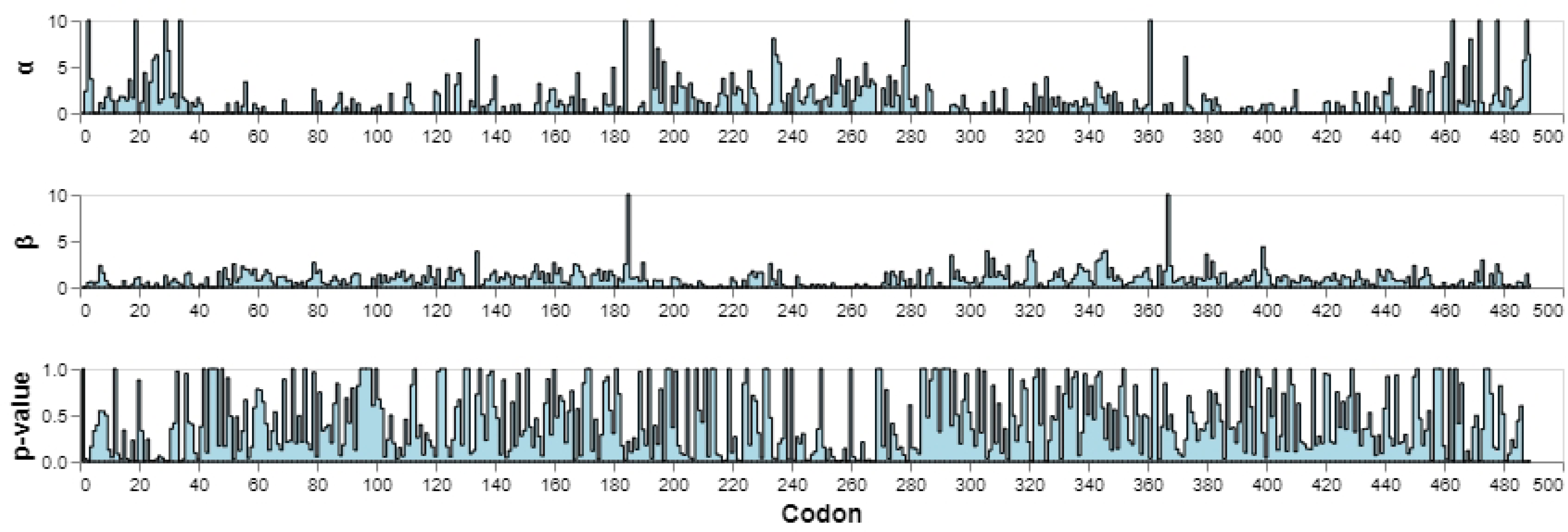
Dense Rate Plot: α; rate of synonymous substitutions at a site, β; rate of non-synonymous substitutions at a site, substitutions in codons at a p-value threshold of ≤0.1. Whereas, X-axis represent the position of the substitution sites within the gene sequence and Y-axis represent the rate of synonymous and non-synonymous substitutions at a site. The p-value indicates statistical significance of these observations.

We visualized the density of synonymous and non-synonymous substitutions across the sequence of E gene. The peaks in the plots indicated regions with α and β substitutions whereas, p-value plot assessed the statistical confidence in these observations. Fig 6 shows the Kernel density estimates of site-level rate estimates in which means are shown with red rules. Estimates above 10 were censored at this value.

**Fig 6.**
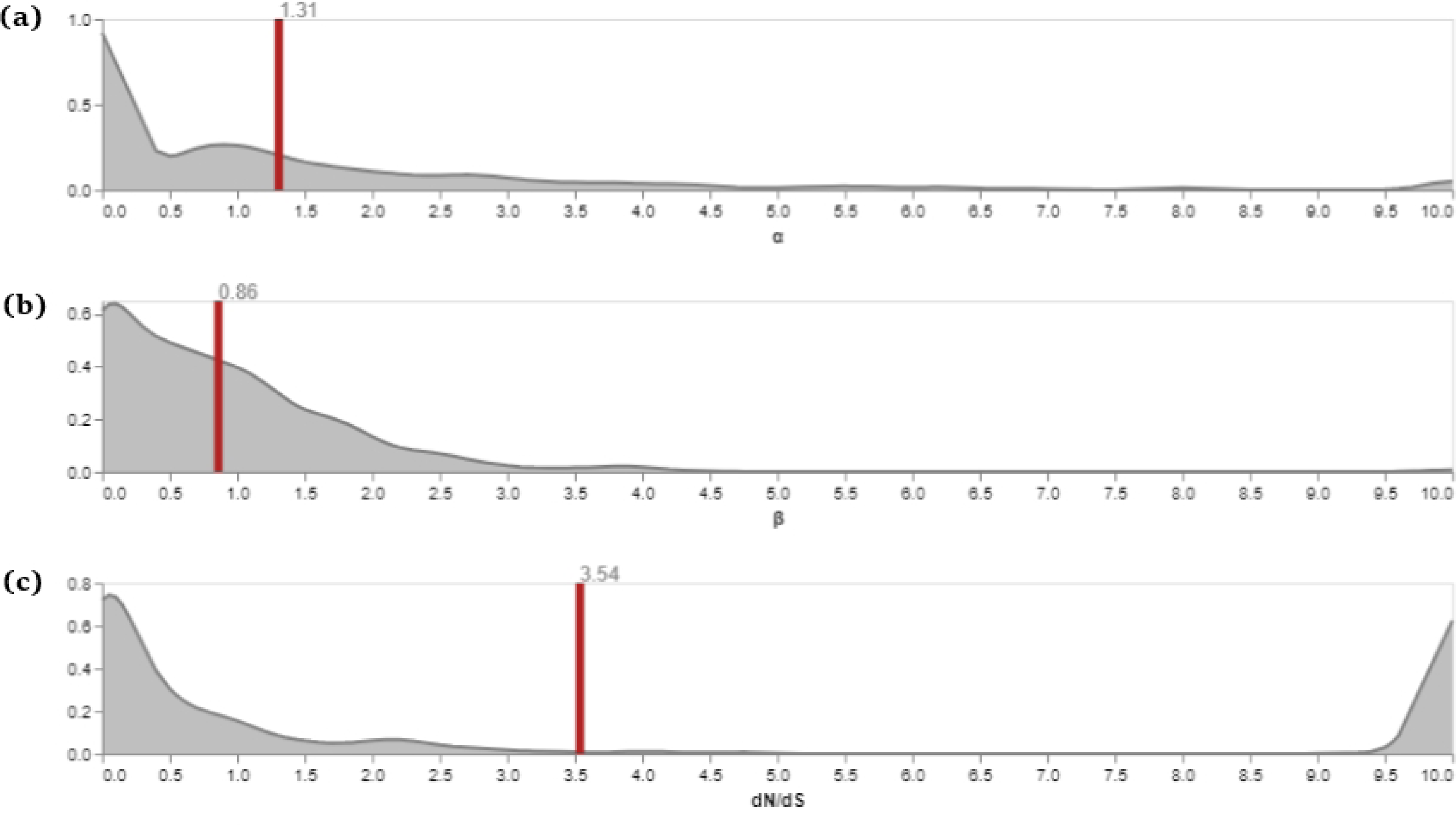
Kernel density estimates of site-level rate estimates of α and β substitutions with overall dN/dS ratio. The plot includes three main components; α and β substitutions along with and dN/dS ratio. X-axis determines site-level rate estimates for substitutions and Y-axis represents the probability density of these rate estimates. The peaks on the Y-axis indicate regions where rate estimates are more concentrated. (a) represents the rate of synonymous substitutions which came out to be 1.31. (b) represents the rate of non-synonymous substitutions which is 0.86. And (c) indicates ratio of overall non-synonymous to synonymous substitutions which is 3.54.

The dN/dS ratio of 3.54 is indicative of some regions in the E gene undergoing adaptive evolution. GARD found evidence of recombinational breakpoints in our data. In this analysis, GARD evaluated a total of 2312 models at a rate of approximately 5.58 models per second. The alignment being examined contained 1404 potential breakpoints which expanded the search space to encompass a staggering 461265714 models each allowing for upto 3 breakpoints to be considered as shown in Fig 7. The presence of recombination breakpoints suggests that genetic exchange or recombination events have likely occurred within the dataset, leading to variations in the genetic sequence.

**Fig 7:**
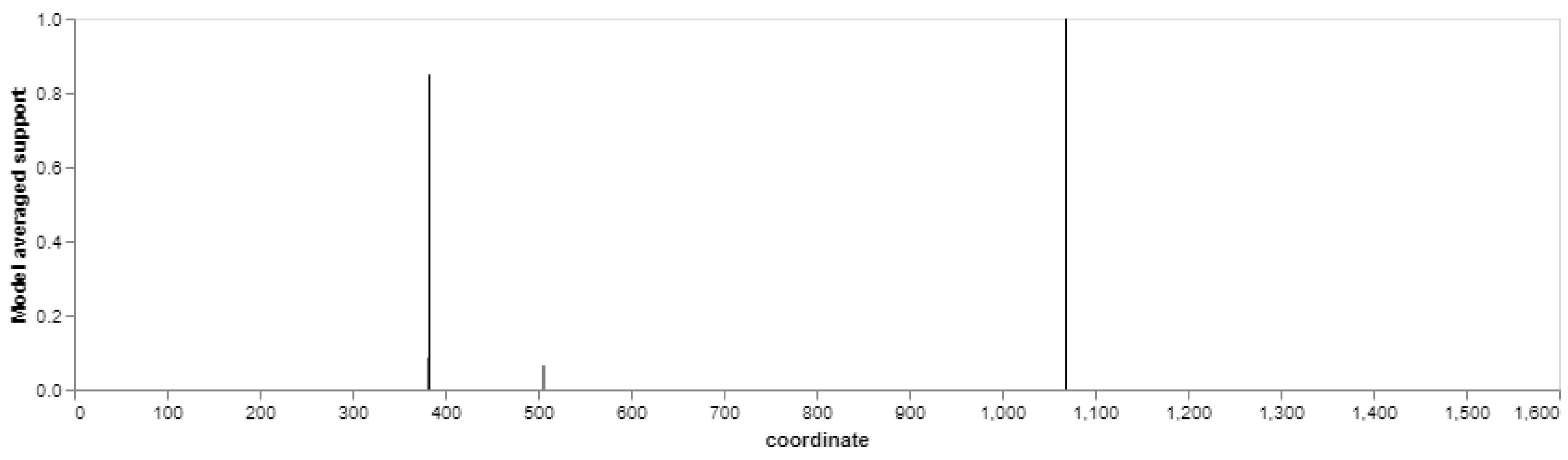
Recombination breakpoints detected in GARD analysis. The X-axis represents the genomic coordinates spanning the range of 0 to 1800, while the Y-axis shows the model average support values ranging from 0.0 to 1.0. Two distinct sharp peaks indicate recombination breakpoints identified at positions 383 and 1069.

The GARD analysis revealed the presence of two prominent sharp peaks at positions 383 and 1069 indicative of recombination breakpoints within the genomic sequence. Fig 8 presents the results of MEME analysis, focusing on the sites within the gene that exhibit evidence of positive selection. Evidence of positive (episodic) selection was detected at 59 specific sites within the gene.

**Fig 8.**
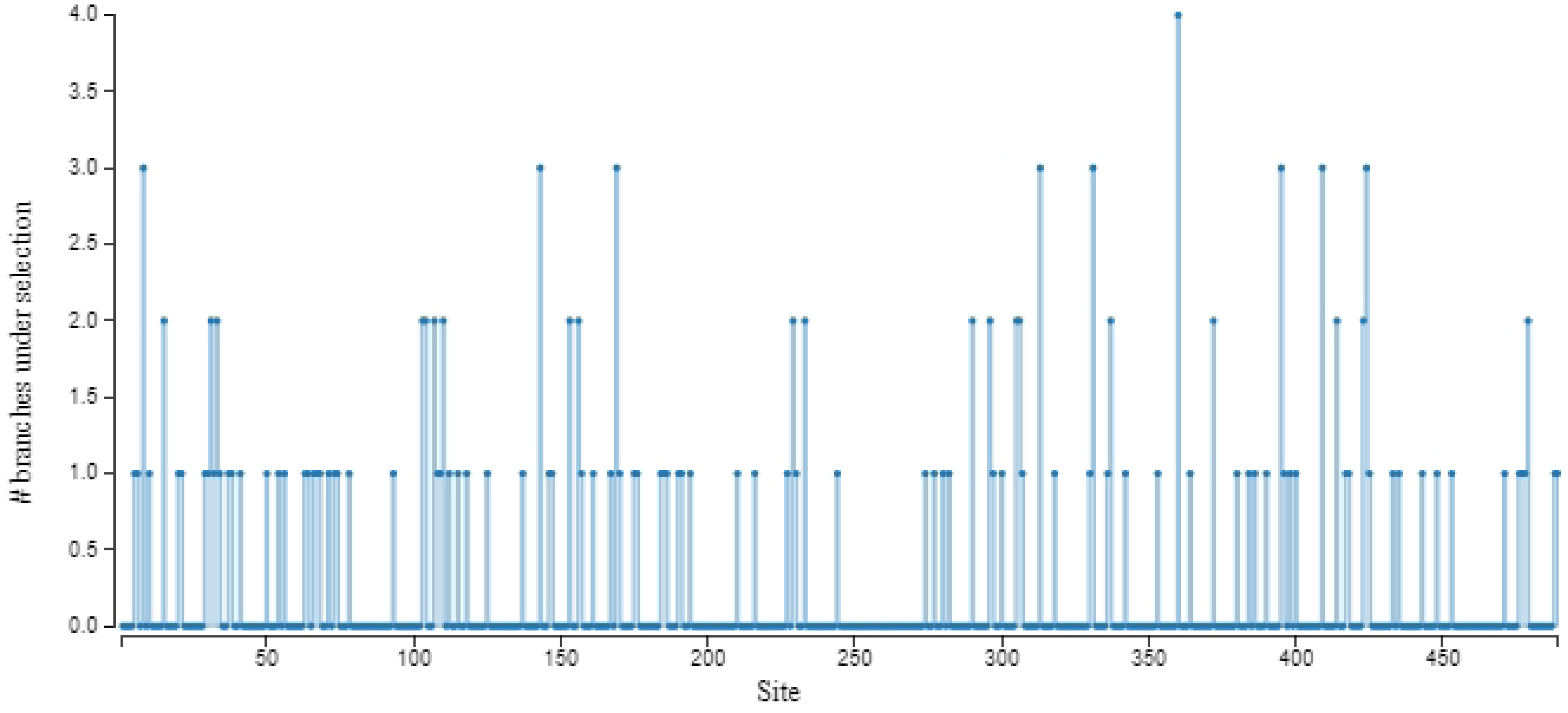
Sites under positive (episodic) selection detected by MEME analysis. The X-axis represents genomic sites within the gene while Y-axis shows the number of branches under positive selection. Several distinct spikes are observed at various sites along the sequence. The spikes indicate specific sites where positive selection has been detected and the height of the spike reflects the strength of evidence for positive selection.

More number of substitutions were observed in the E gene of DENV-1 (11053), DENV-3(11069) and DENV-4(11070) as shown in Fig 8. Whereas, Fig 9 represents the phylogenetic tree generated through MEME analysis, and depicts the evolutionary relationship among the E gene sequences from DENV-1, DENV-2, DENV-3 and DENV-4.

**Fig 9.**
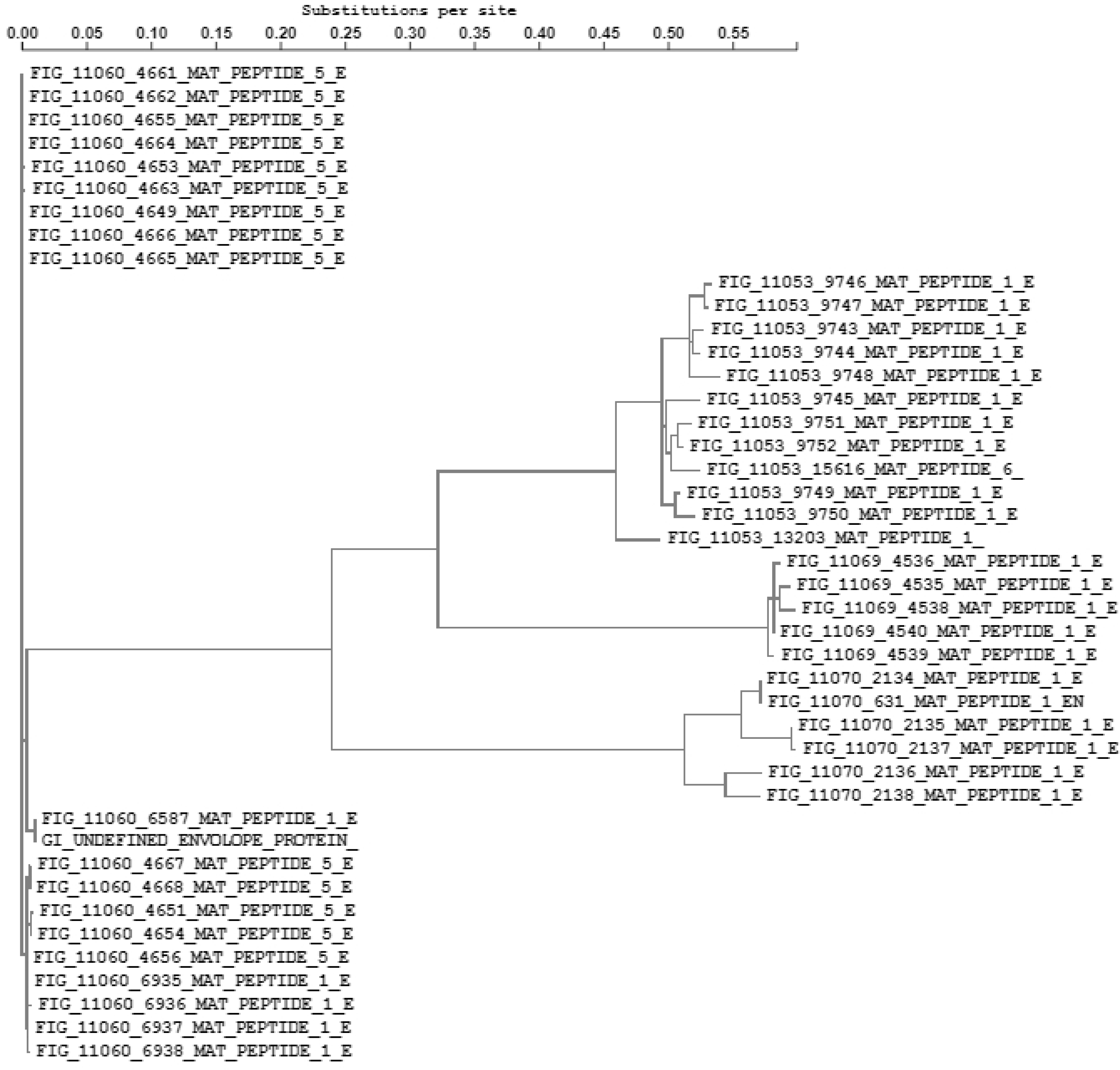
Phylogenetic tree showing substitutions per site for E-gene of DENV serotypes.

The sequences from all four serotypes exhibit a range of substitutions per site with some branches showing values exceeding 0.50 substitutions per site. aBSREL found evidence of episodic positive selection on 12 out of 70 branches in your phylogeny. A total of 70 branches were formally tested for positive selection. Assessed significance using the LRT with a significant threshold at p ≤ 0.05, and the threshold was adjusted to account for multiple testing. The LRT values exceeding this threshold were considered statistically significant, indicating that positive selection is supported at these branches. The positive selection detected with the statistical significance of p ≤ 0.05 indicated that the observed selection at the particular branch is unlikely to have occurred by chance. 12 branches showed LRT values significantly exceeding the threshold set at p ≤ 0.05 even after adjustments for multiple testing. The highest LRT value of 41.2695 observed in one of the DENV-1 E gene as shown in Fig 10.

**Fig 10.**
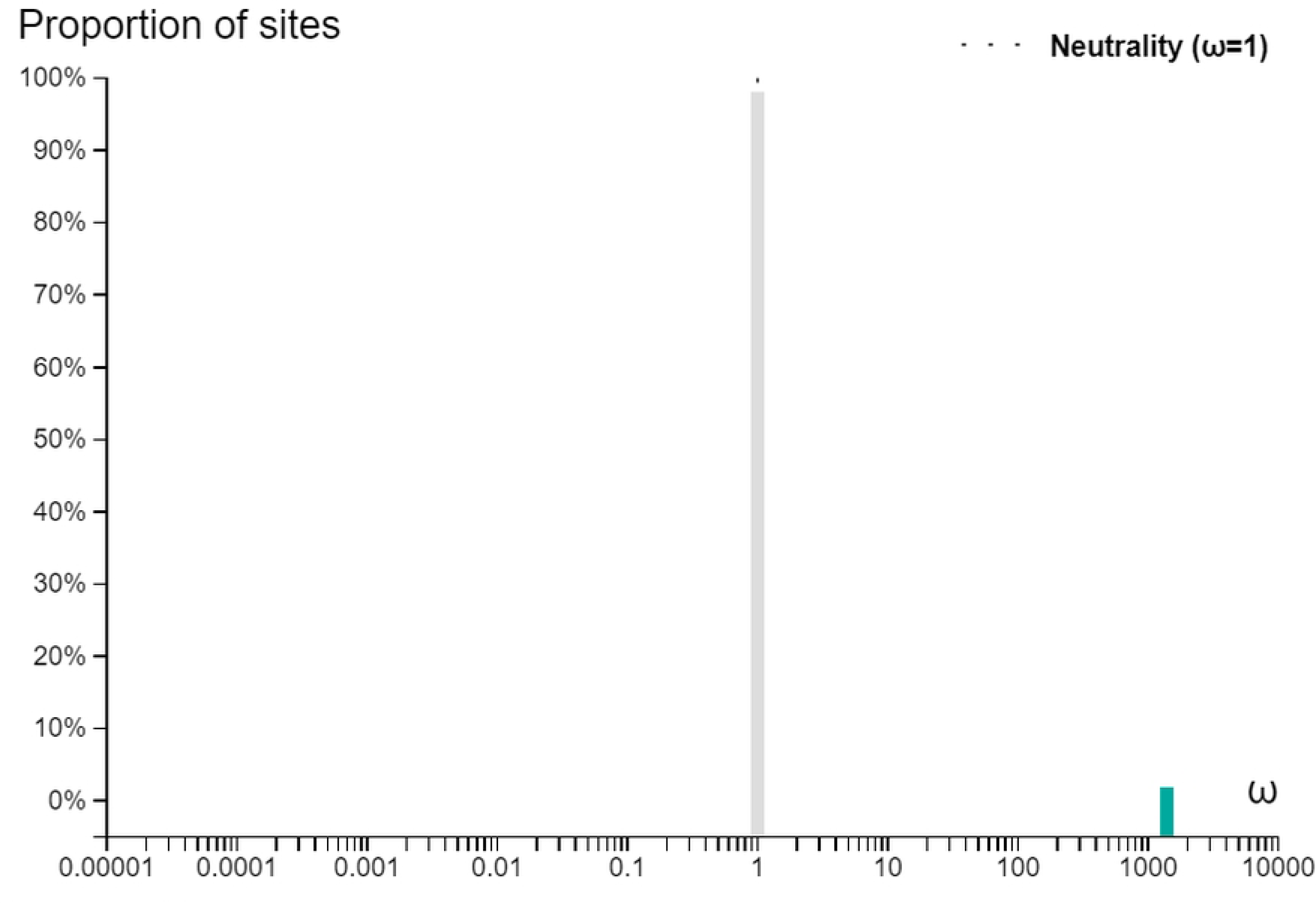
Highest LRT (41.2695) observed in DENV-1 branch signifying the evidence of diversifying selection.

FUBAR identified pervasive diversifying selection at 18 sites and pervasive purifying selection at 50 sites. LRT in BUSTED provided compelling evidence of episodic positive selection within the dataset, with a p-value of 0.000. Fig 11 shows the cumulative distribution of LRT broken down by contribution of individual sites.

**Fig 11.**
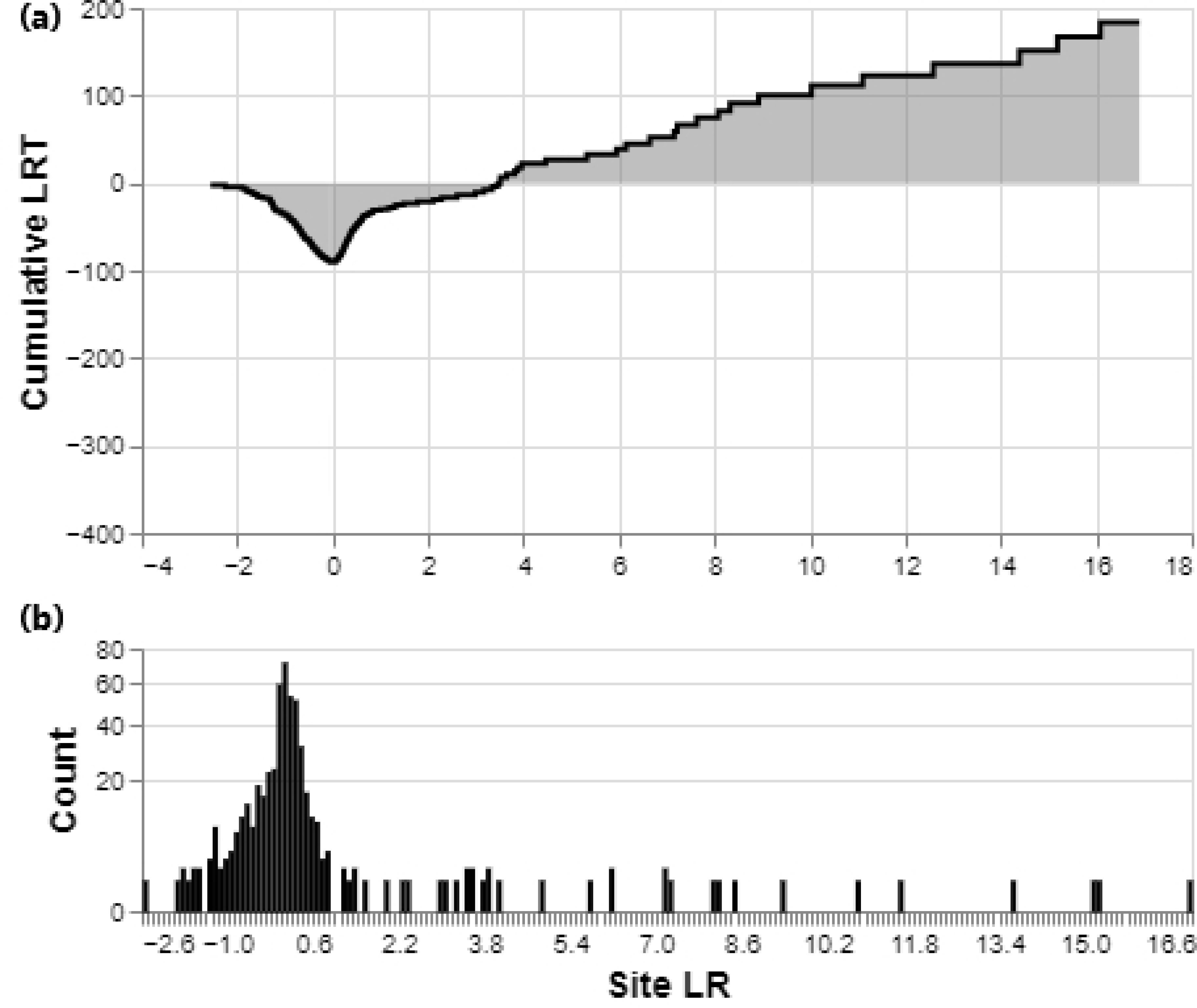
(a) Cumulative distribution of LRT and Site LR. Y-axis representing the cumulative LRT scores and X-axis showing site LRT contribution. (b) Site Count vs. Site LR. The Y-axis represents the count of sites and site LRT values are shown on the X-axis.

There were 25 sites with ER ≥ 10 for positive selection. The downward and upward sloping curve in the graph signifies the cumulative sum of LRT values when we consider more and more sites within the gene. The steep rise in the curve from left to right suggests a cluster of sites with a less negative and more positive selection sites. Fig Contrasting with the aforementioned graph, which showed a cumulative distribution of LRT values illustrating the overall pattern of positive selection, the Fig delves into the diversity of selective pressure acting on different sites providing a more detailed view of the individual site specific LR values.

Moreover, The coexistence of peaks in both the negative and positive LRT value ranges indicated heterogeneity in the data.

### Nucleotide substitution model

When we compared 3H (three-nucleotide substitution) model, 2H (two-nucleotide substitution) model and 1H (one-nucleotide substitution) model, the highest LRT value of 216.703 with p-value of 0.000 obtained for 3H vs. 1H model. For 3H vs. 2H; LRT value was 75.681 with p-value 0.000 and for 2H vs. 1H; LRT value was 141.022 with a p-value of 0.000. The nucleotide substitution pattern, evidence ratio and site log likelihood is shown in Fig 12 (a to c).

**Figure 12.**
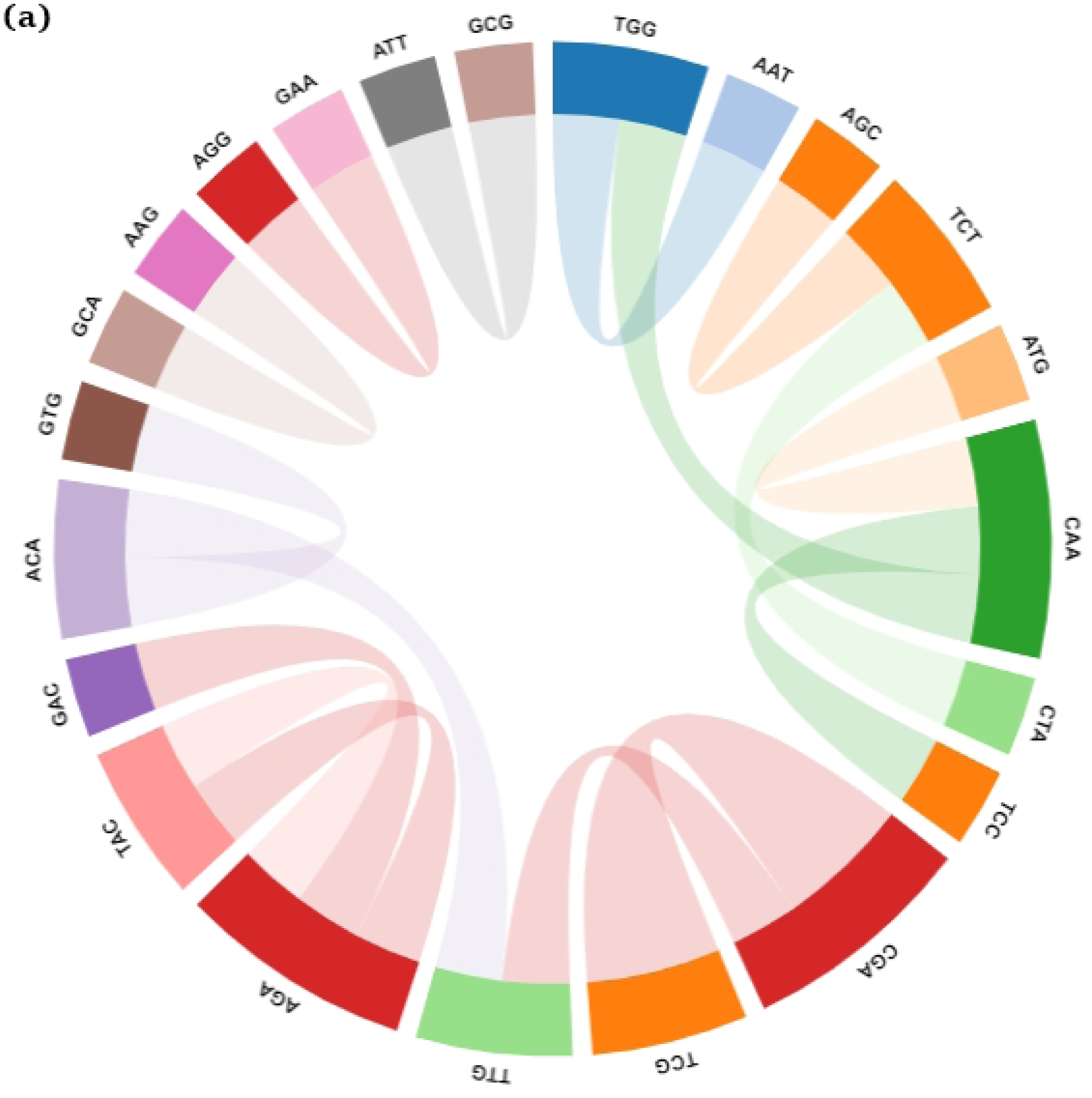
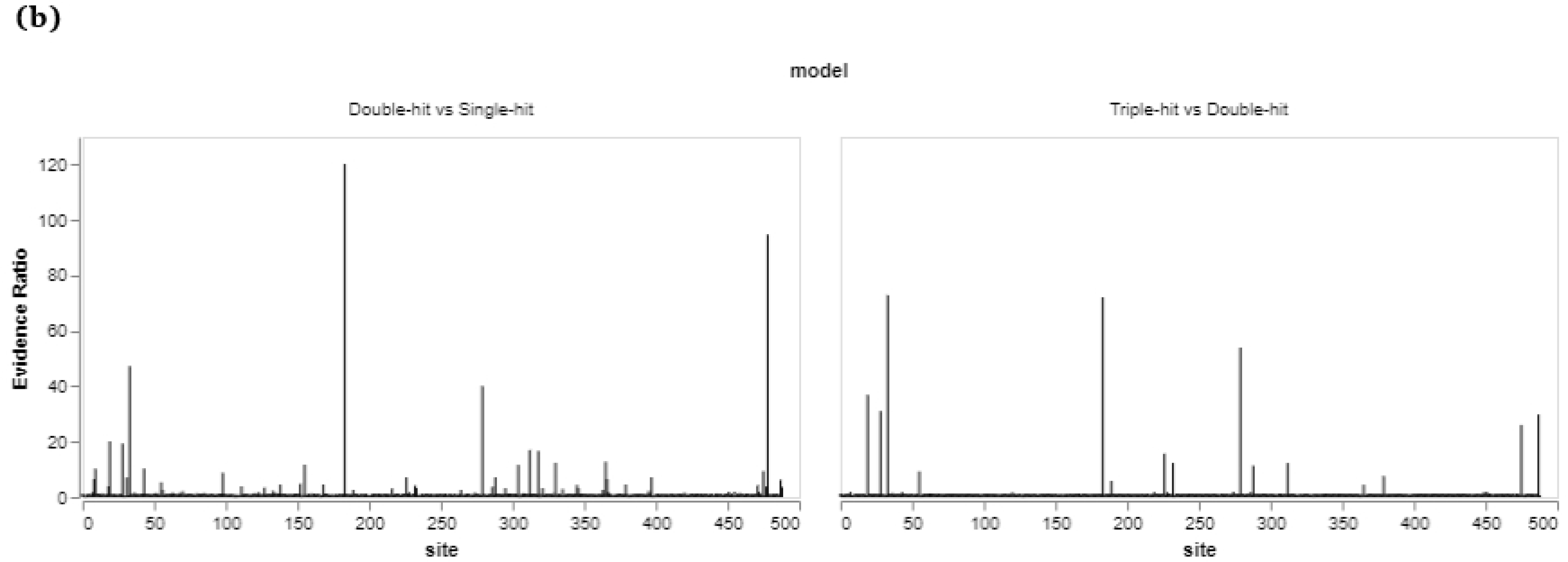
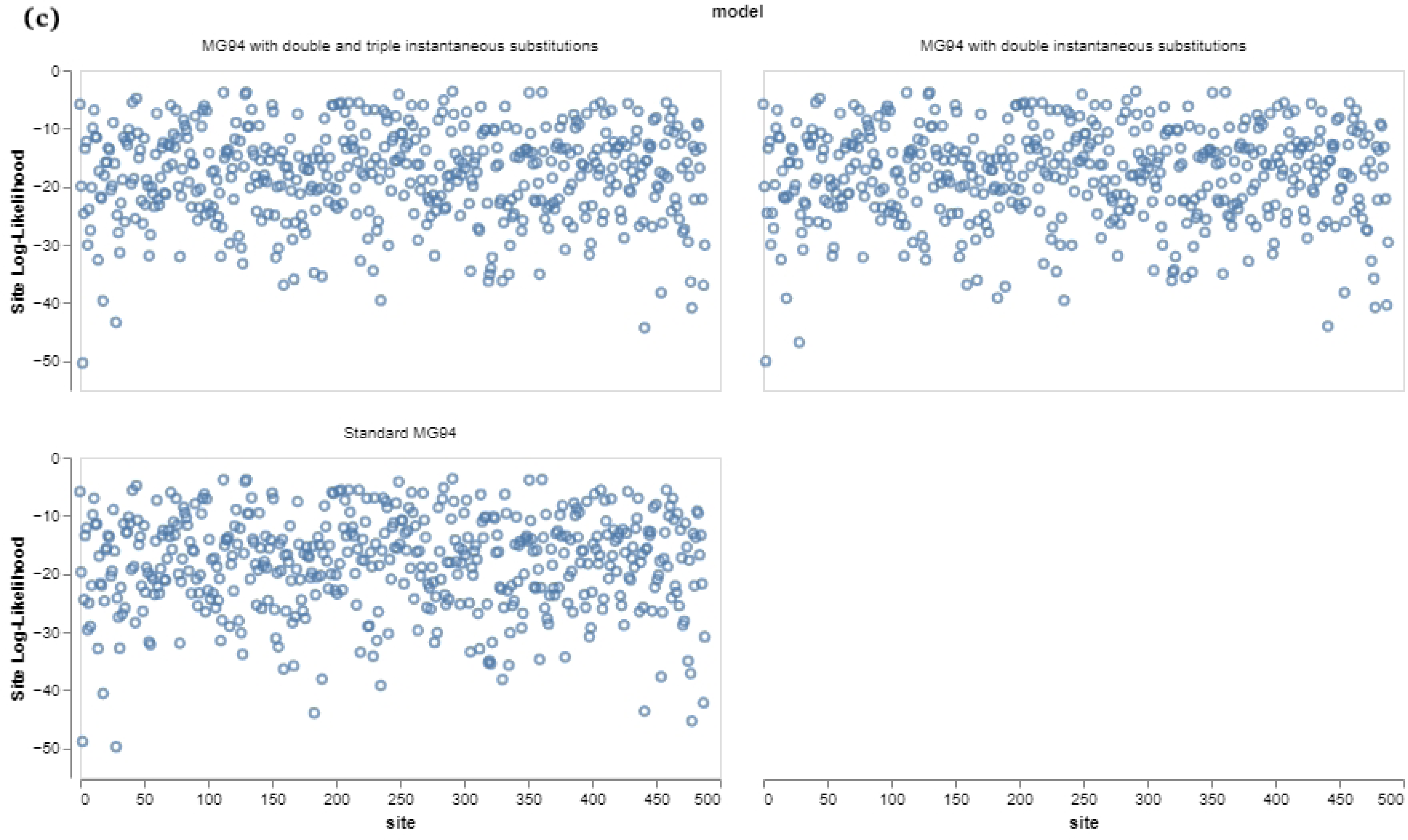
(a) LRT analysis comparing three nucleotide substitution models (3H, 2H and 1H) for site substitutions (b) Evidence Ratio of Site specific nucleotide substitution in double hit vs. single hit and triple hit vs. double hit substitutions analysis (c) Site Log-Likelihood analysis representing site log likelihood on Y-axis and sites on X-axis. The models include MG94 with Double and Triple instantaneous substitutions. MG94 with Double instantaneous substitutions, Standard MG94. Variations in log likelihood values suggest heterogeneous substitution patterns across the gene.

The data strongly support the 3H model (allowing for three-nucleotide substitutions) as the best model for explaining the substitution patterns in your dataset. This suggests that multiple-nucleotide substitutions are occurring in our data, and these substitutions significantly improve the model fit compared to models that allow only one-nucleotide substitutions (1H) or two-nucleotide substitutions (2H).

## Discussion

Dengue infection was believed to be caused by antigenically four diverse serotypes, DENV 1 to 4 sharing over 65% of their genomes. Announcement of a 5th DENV serotype in 2013, has raised a concern in public health departments [3]. To date, no evidence has been found regarding the emergence of DENV-5 in Pakistan. It also raises a speculation that there might be new serotypes which remain undetectable. New serotypes may evolve as a result of genetic bottleneck, natural selection and genetic recombination. Despite being a region with a high occurrence of dengue infections, our understanding of dengue virus evolution has been limited due to the insufficient availability of dengue genomic data from Pakistan. Previous studies on dengue in Pakistan have been primarily focused on localized outbreaks which typically involve single serotype or genotype specific studies from same or closely related strains.

The phylogenetic analysis has revealed a close genetic relationship between DENV-2, the most prevalent and DENV-4, the latest known serotype to have emerged in Pakistan. Despite their distinct serotypes and shared clade, these have significant implications for our understanding of DENV dynamics in the region. The clustering of these two serotypes within the same clade suggests a common evolutionary history and genetic similarity potentially indicating shared ecological niches or environmental conditions conducive to their co-circulation. In Pakistan, current mosquito interventions have proven ineffective in preventing dengue, despite the country being heavily impacted by the disease and contributing significantly to the global dengue burden. Consequently, there is an urgent need for Pakistan to consider the adoption of the dengue vaccine. The shared genetic similarity may necessitate adjustments in vaccine formulations to ensure broader coverage against both serotypes. It could also have implications for cross-immunity and disease severity. If individuals exposed to one serotype develop partial immunity to another due to genetic similarity, it may affect the pattern of disease spread and severity of infection. This information is crucial for healthcare providers and policymakers to anticipate and manage DENV outbreaks [4]. A study was conducted to analyze genetic diversity present within the E gene of DENV-3 serotype so that prevalent amino acid modifications and variability in E protein could be observed [5].

Nucleotide substitution patterns in the genome can be analyzed by Transition/Transversion bias and the R value of 1.94 suggested a moderate bias towards transitions over transversions in the E gene. Similarly, evolutionary analysis of DENV-4 E gene was conducted to observe evolutionary patterns and despite abundant non-synonymus variations, no evidence was found for adaptive evolution in any codon or gene. The combination of a high number of sites (segregating), moderate diversity of nucleotides (π), and a Tajima’s D value greater than zero indicates complex evolutionary dynamics at play in these 43 sequences. These results prompt further investigation into the specific factors driving genetic diversity and the potential role of selection in shaping the genetic landscape of these sequences. Such moderate genetic diversity can be indicative of a complex evolutionary history, including both ancestral genetic variation and more recent evolutionary events [6]. FEL and SLAC determined sites of codons in the E gene alignment data under selection [7]. 59 sites highlighted by FEL provide a valuable insight into how natural selection has shaped genetic diversity at these specific sites within the DENV E gene. aBSREL found 12 branches to be under episodic positive selection. The role of episodic positive selection has also been observed in VP1 and VP3 of foot and mouth disease virus [8]. The results provide valuable insights into the complex forces shaping genetic diversity within the analyzed lineages. Episodic diversifying pressure was also used to find codon sites in ORF of HoBi like pestivirus under selection [9]. The FUBAR analysis has revealed two important aspects of selection within the dataset: Pervasive Positive/Diversifying Selection: evidence of pervasive diversifying selection was found at 18 specific sites. This suggests that these 18 sites have experienced a consistent trend of favoring genetic variations that lead to changes in the amino acid sequence, potentially due to environmental or functional advantages. Such sites may be adapting to new conditions or evolving to perform specific functions more effectively. The analysis identified pervasive negative selection at 50 sites. Purifying selection is often associated with the preservation of critical protein functions or structural integrity. Mutations that negatively impact fitness are removed or suppressed, leading to the persistence of the original genetic code. The LRT comparing the 3H model to the 1H model has a very high LRT value, showing that the 3H model provides a better fit to the data in a significant manner compared to the 1H model [10].

Among RNA viruses, recombination events exhibit varying frequencies in positive sense ssRNA viruses where certain families such as Picornaviridae display elevated rates while others like Flaviviridae experience sporadic occurrences or in case of Leviviridae, virtually none at all. The identification of only two recombination breakpoints in the DENV 1-4 serotypes from Pakistan, raises intriguing questions about the underlying mechanisms driving genetic diversity in these viruses [11]. Flaviviruses are known for their ability to undergo occasional recombinations, and the scarcity of such events in the DENV genome, as observed here, aligns with the virus’s evolutionary strategy. It appears that DENV has developed mechanisms to predominantly rely on mutations generated during replication and interactions with the host immune system, as opposed to frequent recombination events. Recombinational hotspots were also investigated in PSV2020 strain of Picornavirus [12].

The results of our study revealed 18 codons to be under diversifying selection (52,57,107,125,144,156,167,169,175,230,306,313,321,331,337,364,386,400) and 46 codons under purifying selection from DENV (1-4) E gene sequences from Pakistan. (3,11,17,19,22,24,25,26,29,30,35,40,184,193,197,207,212,217,218,222,223,230,234,235,241,24 5,246,252,255,256,257,259,261,262,263,265,267,268,273,279,326,456,463,467,469,472). Those codons were considered that were confirmed by at least two selection pressure analysis methods. More no. of purifying codons also supports the results of Tajima’s D test, indicating the DENV genome experiencing balancing selection. Similarly, a major role of negative selection was also observed in the HoBi like pestivirus genome in shaping its evolution [9]. Remarkable heterogeneity was observed across different sites within the E gene indicating some regions under strong purifying selection to maintain function while others may be under positive selection to adapt to changing environments.

However, the study has certain limitations, such as the absence of DENV serotypes sequencing data from Pakistan collected in 2023 and the limited availability of complete genome DENV-4 sequence from Pakistan. Having a more extensive dataset of complete genome sequences would have provided a more comprehensive understanding of the situation regarding dengue virus in Pakistan. Evolutionary analysis of the E gene and complete genome of DENV serotypes consistently revealed a close evolutionary relationship between DENV-1 and DENV-3, forming a distinct clade. DENV-2 and DENV-4 are more genetically divergent from the other two serotypes with DENV-4 being the most genetically distinct. This newfound understanding of genetic relationships can be utilized to develop predictive models for DENV outbreaks in Pakistan encompassing both genetic and environmental factors thus optimizing preparedness and response strategies. Models can also be developed to predict new evolving serotypes from genomic data of available serotypes.

## Materials and methods

### Data collection

#### Dataset 1

Retrieved 448 sequences of Pakistani Dengue virus isolates from NCBI and ViPR database [13]. The sequences encompassed a range of genetic data, comprising complete genome sequences. We exclusively considered sequences that came with associated location information and collection dates. Among 448 sequences, the repeated sequences with same accession no. and size less than 500 bp were removed. Most of the sequences were collected over a span of years from 1994 to 2023 and a total of 43 complete genome sequences were further analyzed. DENV-1 (7 sequences), DENV-2 (24 sequences), DENV-3 (10 sequences) and DENV-4 (2 sequences). Data was composed of all four serotypes. Reference sequences for all four serotypes were included in the study. Due to unavailability of current serotypes data from 2023, 6 recently reported sequences were included from outside Pakistan to have a clear picture of recent mutational patterns in dengue serotypes. After preparing four separate files containing serotype specific data, concatenated all files for Multiple Sequence Alignment (MSA). MUSCLE alignment was run on MEGA-X [14].

#### Dataset 2

We acquired all Pakistani dengue virus E gene nucleotide sequences documented till now from the ViPR database. This dataset encompassed 46 E protein sequences from DENV-1 to DENV-4.

### MSA and phylogenetic analysis

Dataset 1 comprising a total 43 complete genome sequences of all four serotypes from Pakistan reported till date along with four reference sequences of dengue virus and 6 sequences other than from Pakistan were retrieved from the same databases mentioned earlier to have a clear understanding of mutational patterns in E gene (data available in supplementary file). The MUSCLE results revealed that every sequence from one serotype had a high degree of similarity to reference sequences of the same serotype. Furthermore, conserved and non-conserved regions in the entire genome were also manually screened. Sequence alignment was also performed for the E gene using the same software.

#### Tamura-Nei model and Maximum Likelihood (ML)

Model for tree construction in MEGA-X was selected by Akaike information criterion (AIC) [14]. The Tamura-Nei model and Maximum Likelihood (ML) approach was used to infer evolutionary history [15]. Bootstrap consensus tree generated from 1000 replicates was used to analyze evolutionary history of taxa. And the branches associated with partitions that had less than 50% of the bootstrap replicates collapsed. In the Bootstrap test with 1000 replicates, the proportion of replicated trees in which linked taxa clustered together was displayed next to branches [16]. The initial tree for heuristic search was created by automatically applying Neighbor-join and BioNJ algorithms to a matrix of pairwise distance calculated using Tamura-Nei model and then choosing the topology with highest log likelihood value. 43 nucleotide sequences were included for the complete genome phylogenetic analysis. The final dataset contained a total of 10817 positions. Phylogenetic analysis was also performed to see evolutionary linkages between E genes of DENV serotypes utilizing the same strategy.

### Statistical analysis to predict Neutrality

Complete genome sequences were used to run Fisher’s exact test to calculate chances of evidence that the serotypes are evolving in a way that’s not random. For each pair of sequences in the datasets A, we looked at the chances of rejecting the idea that they are evolving neutrally and favoring the idea that they are being positively selected. p-value < 0.05 was significant. The variances were estimated by the bootstrap method.

The codon-based Z test was utilized to assess neutrality between sequences with significance considered when P values were less than 0.05 (at the 5% level). This test relies on dS and dN to refer to the number of synonymous and non-synonymous substitutions per site. To analyze evolutionary patterns. The Nei-Gojobori method [17]. was employed for these analyses, and pairwise deletion option was used to remove ambiguous positions for each sequence pair, yielding a final dataset with 3275 positions.

To estimate Gamma parameters for site rates, we utilized discrete Gamma distribution with a shape parameter of 0.3926. Tamura-Nei model featuring a gamma distribution was used to determine the patterns and rates of substitution (+G) [18]. The gamma distribution comprising 5 categories were incorporated to account for variations in evolutionary rates among sites. The estimated ML values and the automatically computed tree topology produced a maximum-log likelihood of −65617.091. We analyzed the position of every codon including first, second, third and non-coding ones, yielding a final dataset of 10,817 positions.

Tajima’s D is a genetic statistical test used in population genetics. It compares genetic diversity in nucleotide sequences to distinguish between random evolution with neutral mutations and non-random evolution influenced by factors like natural selection or changes in population size. Neutral mutations have no effect on an organism’s survival, while non-neutral mutations are influenced by selection. In this analysis, a set of 43 nucleotide sequences was examined.

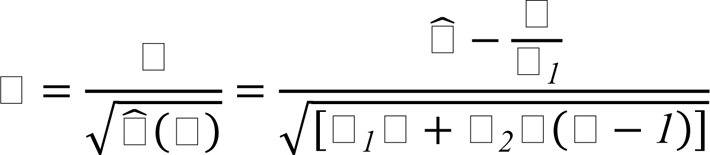

The analysis considered all codon positions, including the 1st, 2nd, 3rd, and noncoding positions. For each set of sequences, the ambiguous positions were eliminated using paired deletion to assure accuracy of results. The total number of positions in the final dataset were 10,817 and the evolutionary analysis was performed on MEGA 11. Mathematical representation of the test is given below.

### Inter-sequence Substitution patterns

To determine if the entire genome sequences were evolving with a similar pattern of substitution or showed noticeable variances in base composition biases, the Disparity index test was used to test the homogeneity of substitutions among sequences. In order to access patterns of substitutions, the non-coding positions first, second, third and non-coding were taken into account resulting in a dataset of 10,817 positions. To access p-values, a Monte Carlo test with 500 repeats was used, with results below 0.05 were of significance. Tamura-Nei (1993) model was also utilized under ML estimate of substitutions matrix to estimate substitution probabilities (r) between nucleotide bases. Where,

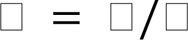

s determine the number of transitional and v determine the number of transversional substitutions per site.

The probabilities were categorized into transitional and transversional substitutions (data available in supplementary file) to ensure relative proportions the sum of substitution probabilities equaled 100% and the determined frequency of nucleotides were A = 32.48%, T/U = 21.19%, C = 20.61%, and G = 25.71%. The Transition/Transversion bias (R) was assessed using the Kimura (1980) 2-parameter model. The model was applied to estimate the probability of substitution (r) between different nucleotide bases differentiating between transitional and transversional substitutions.

### Selective pressure analysis

The ratio of dN to dS in DENV E-gene sequences were determined using different statistical methods; SLAC, FEL, MEME and FUBAR. We utilized SLAC and FEL both implemented in the HyPhy package, and the sites with p-value <0.1 were significantly considered. For FEL the statistical significance was determined by assessing the results against the asymptotic χ2 (chi-squared) distribution. This analysis considered variations in synonymous (silent) mutation rates between different sites in the dataset. The Nucleotide GTR model was selected for analysis. FEL is often used to identify sites in a genetic sequence that may be evolving under positive or negative selection. MEME analysis was performed to find the diversifying selection at various points during evolutionary history. The statistical significance of this discovery was based on a p-value threshold of 0.1. We conducted MEME analysis to find the diversifying selection at various points during evolutionary history. An evolutionary model based on LRT; Likelihood Ratio Test was used for analysis. At particular codons, we searched for instances of positive selection considering it significant when p-value was less than 0.1. We used the Bayes empirical Bayes approach to identify lineages where diversifying selection took place at a specific codon. Branch types that had codons with Bayes factor greater than 1 were known to be subject to episodic diversifying selection. The dN/dS ratio was divided into three groups for BSR analysis using the MG94 model. If the corrected p-value dropped below 0.1, the null model was deemed to be rejected using the BSR approach. For aBSREL, 70 branches were explicitly evaluated for diversifying selection. The LRT was used to determine significance with a p-value threshold of 0.05. FUBAR in the HyPhy package was used to find purifying and diversifying selection sites with a subsequent likelihood of 0.9. BUSTED v4.1 analysis was performed on the E gene alignment using HyPhy (v2.5.48) integrated in datamonkey. The analysis included site to site synonymous rate variation by setting evidence ratio threshold 10. Using the HyPhy package, a multi-hit analysis was performed to determine the potential impact of additional base hits in our data.

## Supporting information

**S1 Table.** Detailed information of complete genome DENV sequences (Dataset 1)

**S1 Figure.** Evolutionary relatedness of dengue virus serotypes and prevalence of genotypes in Pakistan.

**S2 Table.** Maximum Likelihood Estimate of Substitution Matrix

**S3 Table.** Maximum Composite Likelihood Estimate of the Pattern of Nucleotide Substitution

**S4 Table.** Maximum Composite Likelihood Estimate of the Pattern of Nucleotide Substitution (transition/transversion)

**S5 Table.** Fixed Effect Likelihood estimates of diversifying and purifying codons

## References

1. Islam A, Deeba F, Tarai B, Gupta E, Naqvi IH, Abdullah M, Dohare R, Ahmed A, Almajhdi FN, Hussain T, Parveen S. Global and local evolutionary dynamics of Dengue virus serotypes 1, 3, and 4. Epidemiology & Infection. 2023 Jan;151:e127.

2. Erb SM, Butrapet S, Roehrig JT, Huang CY, Blair CD. Genetic Adaptation by Dengue Virus Serotype 2 to Enhance Infection of Aedes aegypti Mosquito Midguts. Viruses. 2022 Jul 19;14(7):1569.

3. Mustafa MS, Rasotgi V, Jain S, Gupta VJ. Discovery of fifth serotype of dengue virus (DENV-5): A new public health dilemma in dengue control. Medical journal armed forces India. 2015 Jan 1;71(1):67–70.

4. Forni D, Sironi M, Cagliani R. Evolutionary history of type II transmembrane serine proteases involved in viral priming. Human Genetics. 2022 Nov;141(11):1705–22.

5. Amarilla AA, de Almeida FT, Jorge DM, Alfonso HL, de Castro-Jorge LA, Nogueira NA, Figueiredo LT, Aquino VH. Genetic diversity of the E protein of dengue type 3 virus. Virology journal. 2009 Dec;6:1–3.

6. Forni D, Sironi M, Cagliani R. Evolutionary history of type II transmembrane serine proteases involved in viral priming. Human Genetics. 2022 Nov;141(11):1705–22.

7. Jagtap S, Pattabiraman C, Sankaradoss A, Krishna S, Roy R. Evolutionary dynamics of dengue virus in India. PLoS pathogens. 2023 Apr 3;19(4):e1010862.

8. Subramaniam S, Mohapatra JK, Das B, Sharma GK, Biswal JK, Mahajan S, Misri J, Dash BB, Pattnaik B. Capsid coding region diversity of re-emerging lineage C foot-and-mouth disease virus serotype Asia1 from India. Archives of virology. 2015 Jul;160:1751–9.

9. Kalaiyarasu S, Mishra N, Subramaniam S, Moorthy D, Sudhakar SB, Singh VP, Sanyal A. Whole-Genome-Sequence-Based Evolutionary Analyses of HoBi-like Pestiviruses Reveal Insights into Their Origin and Evolutionary History. Viruses. 2023 Mar 11;15(3):733.

10. Zhang Q, Liu J, Han S, Wang B, Su Q, Yuan G, He H. Genetic and evolutionary characterization of avian paramyxovirus type 4 in China. Infection, Genetics and Evolution. 2021 Jul 1;91:104777.

11. Ochwo-Ssemakula M, Mbewe W, Badji A. Computational evidence that the Ugandan Passiflora virus likely evolved from the Bean common mosaic necrosis virus primarily through recombination.

12. Chen QY, Sun ZH, Che YL, Chen RJ, Wu XM, Wu RJ, Wang LB, Zhou LJ. High Prevalence, Genetic Diversity, and Recombination of Porcine Sapelovirus in Pig Farms in Fujian, Southern China. Viruses. 2023 Aug 17;15(8):1751.

13. Pickett BE, Sadat EL, Zhang Y, Noronha JM, Squires RB, Hunt V, Liu M, Kumar S, Zaremba S, Gu Z, Zhou L. ViPR: an open bioinformatics database and analysis resource for virology research. Nucleic acids research. 2012 Jan 1;40(D1):D593–8.

14. Tamura K, Stecher G, Kumar S. MEGA11: molecular evolutionary genetics analysis version 11. Molecular biology and evolution. 2021 Jul 1;38(7):3022–7.

15. Tajima F. Statistical method for testing the neutral mutation hypothesis by DNA polymorphism. Genetics. 1989 Nov 1;123(3):585–95.

16. Forni D, Sironi M, Cagliani R. Evolutionary history of type II transmembrane serine proteases involved in viral priming. Human Genetics. 2022 Nov;141(11):1705–22.

17. Nei M, Gojobori T. Simple methods for estimating the numbers of synonymous and nonsynonymous nucleotide substitutions. Molecular biology and evolution. 1986 Sep 1;3(5):418–26.

18. Tamura K, Nei M. Estimation of the number of nucleotide substitutions in the control region of mitochondrial DNA in humans and chimpanzees. Molecular biology and evolution. 1993 May 1;10(3):512–26.

